# Endogenous gene tagging in the model brown alga *Ectocarpus* using a simplified CRISPR/Cas method

**DOI:** 10.64898/2026.08.21.745938

**Authors:** Alexandre Paix, Morgane Raphalen, Elena Avdievich, Ferran Agullo, Remy Luthringer, Susana M. Coelho

## Abstract

Brown algae represent one of the few eukaryotic lineages to have independently evolved complex multicellularity, providing a powerful comparative system for investigating the molecular and evolutionary principles underlying multicellular development. *Ectocarpus* has emerged as the principal model for this lineage, supported by extensive genomic and transcriptomic resources. However, mechanistic and functional studies have remained limited by the available reverse-genetic tools. While recent CRISPR-Cas developments have enabled targeted gene knock-outs, the lack of knock-in (KI) approaches for endogenous protein tagging and precise genomic insertion remains a major experimental bottleneck. Here, we establish a comprehensive CRISPR-Cas genome-engineering framework for *Ectocarpus* that enables both targeted gene disruption and precise genomic insertion. We demonstrate efficient knock-in of multiple peptide tags at endogenous loci, enabling direct analysis of native proteins. By combining robust gene knock-out with endogenous protein tagging, this framework substantially expands the experimental possibilities for brown algal research and establishes *Ectocarpus* as a genetically tractable system for functional genomics, providing a foundation for genome engineering across stramenopiles.

## INTRODUCTION

Brown algae (Phaeophyceae) are amongst the most ecologically important and evolutionarily distinctive multicellular organisms on Earth (Bringloe et al., 2020). As dominant habitat-forming species in many coastal ecosystems, they underpin marine biodiversity and contribute substantially to global carbon cycling (Bringloe et al., 2020; Brodie et al., 2017). Brown algae also occupy a unique phylogenetic position within the Stramenopiles, having evolved complex multicellularity independently from both animals and land plants more than one billion years ago (Bringloe et al., 2020; Cock et al., 2010; Coelho, 2024). This independent evolutionary origin makes them particularly valuable for investigating fundamental questions concerning the evolution of multicellular development, cellular differentiation, life cycles, sex determination, and genome evolution (Barrera-Redondo et al., 2025; Batista et al., 2024; Cock et al., 2014; Coelho, 2024; Coelho et al., 2011; Coelho et al., 2020; Lotharukpong et al., 2026).

Among brown algae, *Ectocarpus* has emerged as the principal model system for molecular, developmental, and evolutionary studies (Coelho, 2024; Coelho et al., 2020). The availability of a complete genome sequence, extensive transcriptomic resources and genomic datasets has enabled important advances in our understanding of developmental regulation in this group of organisms (Bourdareau et al., 2021; Gueno et al., 2022; Liu et al., 2024; Vigneau et al., 2026). These resources have established *Ectocarpus* as a powerful system for comparative biology and evolutionary developmental genetics.

Despite this progress, the ability to directly test gene function has remained a major limitation. Functional studies in *Ectocarpus* have historically relied on forward genetic screens and comparative expression analyses (Coelho et al., 2011; Godfroy et al., 2017; Godfroy et al., 2023; Macaisne et al., 2017). While these approaches have generated numerous hypotheses regarding the genetic basis of developmental and evolutionary processes, the lack of efficient reverse genetic tools has constrained mechanistic investigation of gene function and cellular processes. The recent introduction of CRISPR-Cas-mediated genome editing in *Ectocarpus* represented an important step toward overcoming this limitation, enabling efficient targeted gene knock-out (KO) and establishing the first reverse-genetic approaches in a brown alga (Badis et al., 2021; Martinho et al., 2025). These advances transformed the study of gene function in this lineage and enable direct testing of developmental regulators and candidate genes. However, these approaches have remained so far restricted to gene disruption, leaving many of the most powerful applications of modern genome engineering inaccessible.

In established model organisms, targeted genomic insertions have revolutionized functional biology (Adli, 2018; Van Vu et al., 2025; Zirin et al., 2022). Knock-In (KI) approaches allow endogenous protein tagging, conditional allele construction, protein localization studies and interaction analyses, and precise manipulation of regulatory sequences. In particular, endogenous tagging has become indispensable for cell biology because it enables visualization of proteins under native regulatory control while avoiding artefacts associated with transgene overexpression (Cho et al., 2022; Kim et al., 2024). Such approaches have transformed our understanding of cellular organization and developmental dynamics across animals, fungi, and plants.

The absence of KI strategies currently represents a major bottleneck for brown algal research. Many outstanding questions in *Ectocarpus* biology, including the cellular mechanisms underlying developmental patterning, life-cycle regulation, sex determination, chromosome dynamics, and host-virus interactions, require the ability to manipulate genes within their native genomic context and to visualize proteins in cells. More generally, the lack of targeted insertion methods has limited the transition of *Ectocarpus* from an emerging model organism to a fully genetically tractable experimental system.

Here, we establish a CRISPR-Cas genome engineering approach for *Ectocarpus* that enables both targeted gene disruption and genomic insertion, and demonstrate knock-in of several peptide tags at endogenous loci. By combining reliable gene knockout with endogenous protein tagging, this approach substantially expands the experimental toolkit for brown algal research, bringing *Ectocarpus* to a new level of genetic tractability and enabling mechanistic studies in a lineage that evolved complex multicellularity independently of plants and animals. More broadly, it provides a foundation for functional genomics across brown algae and establishes a genome engineering framework that can accelerate research throughout the stramenopile lineage.

## RESULTS

### Transformation optimization

To maximize the efficiency of subsequent knock-in (KI) experiments, we first optimized our previously established CRISPR-Cas KO workflow. *Ectocarpus* gametophytes release tens of thousands of haploid gametes in response to conditions mimicking low tide (Coelho, 2024; Coelho et al., 2011; Coelho et al., 2012). A fraction of these gametes settle and undergo asexual development into partheno-sporophytes, which subsequently produce gametophytes through spores (Coelho et al., 2012). This provides a large population of haploid cells that can be directly subjected to genome editing and screened for edited individuals.

We used polyethylene glycol (PEG)-mediated delivery of Cas ribonucleoprotein (RNP) complexes, an approach successfully applied in several algal species to facilitate cellular uptake of RNPs (Blomme et al., 2021; Martinho et al., 2025), presumably through PEG-induced precipitation and changes in membrane permeability(Maas and Werr, 1989). To optimize editing conditions, we targeted *APT*, encoding adenine phosphoribosyltransferase, which provides a convenient selectable readout for gene disruption. APT converts adenine to AMP and also metabolizes the toxic adenine analogue 2-fluoroadenine (2-FA). Consequently, wild-type (WT) cells are sensitive to 2-FA, whereas loss-of-function *APT* mutants are resistant, allowing edited individuals to be directly selected and making *APT* a useful marker for both optimization of genome editing and subsequent co-transformation experiments (**Figure S1A**).

In *Ectocarpus*, gametes can be transformed by PEG during the first 2 h following their release, before cell wall formation begins (Martinho et al., 2025). We therefore first tested whether shortening the interval between gamete release and transformation could improve editing efficiency, using Cas9 or Cas12a RNPs targeting *APT* and the recovery of 2-FA-resistant individuals as a quantitative readout. In the standard protocol (Martinho et al., 2025), large numbers of gametophytes are concentrated and induced to release gametes into a large volume of seawater (SW), followed by gentle centrifugation to concentrate the released gametes. Because these steps are time-consuming, transformation is often performed close to the end of the 2 h competence window. We reasoned that inducing gamete release in a small volume of SW and directly recovering the released gametes would eliminate the centrifugation steps and substantially shorten the procedure. Using this simplified protocol, we were able to perform transformation as early as 35 min after induction of gamete release (**Figure S1B**). Importantly, early transformation produced more 2-FA-resistant *APT* mutants than transformation performed 1h45min after release. Thus, in addition to simplifying the procedure, minimizing the time between gamete release and RNP delivery markedly improved transformation efficiency.

To further streamline the protocol, we tested whether altering the ratio of components in the transformation mixture could improve editing efficiency. Previously, 100 µL of gametes in SW were mixed with 20 µL of Cas9 or Cas12a RNPs and 120 µL of PEG solution. We compared this with a reduced-volume protocol using only 20 µL of gametes, while maintaining the same volumes of RNP and PEG solutions, thereby increasing their concentrations relative to the number of gametes. Depending on the experiment, this modification either maintained or increased editing efficiency (**Figure S1C**), despite using substantially fewer gametes. The reduced gamete requirement offers several practical advantages: fewer gametophytes are needed for each transformation, more conditions can be tested in parallel, and a larger number of independent transformations can be performed from a single gamete release.

We next compared the editing outcomes generated by Cas9 and Cas12a to select the most suitable nuclease for KI. Cas9 generates a double-strand break (DSB) close to the guide RNA PAM (protospacer adjacent motif), producing predominantly blunt DNA ends, whereas Cas12a cleaves the two DNA strands at staggered positions distal to the PAM, generating 5′ overhangs. Sequencing of multiple 2-FA-resistant *APT* alleles revealed that Cas9 generally produced smaller insertions and deletions (InDels) than Cas12a (**Figure S1D–F**). We therefore selected Cas9 for subsequent KI experiments, reasoning that its more localized editing outcomes would minimize unwanted sequence alterations at the targeted locus. Although Cas9 generated fewer 2-FA-resistant individuals overall than Cas12a, the optimized protocol nevertheless yielded sufficient numbers of Cas9-edited individuals for downstream KI experiments (**Figure S1E**).

### Establishing targeted knock-in at the *APT* locus

We next used the *APT* locus as a selectable system to establish conditions for Cas9-mediated KI. As donor templates, we designed single-stranded oligodeoxynucleotides (ssODNs), which are widely used for targeted sequence insertion in genome editing (Dewari et al., 2018; Jin et al., 2025). The ssODNs encoded an HA epitope tag (27 bp), followed by a STOP codon and an additional nucleotide designed to disrupt the *APT* coding sequence upon integration. Thus, successful insertion, like a KO-inducing InDel, was expected to confer resistance to 2-FA. The HA sequence additionally provided a readily detectable signature of successful KI that could be identified by PCR genotyping and confirmed by sequencing.

We reasoned that, if *Ectocarpus* possesses sufficient endogenous DNA ligation activity, providing donor DNA at high concentration could promote its capture at Cas9-induced DSBs through non-homologous end joining (NHEJ), effectively joining the chromosomal ends through the donor sequence. We therefore tested ssODNs in both sense (S) and antisense (AS) orientations, defined relative to the *APT* coding sequence, as well as a mixture containing both oligo orientations in the same transformation (**Figure 1A**).

**Figure 1.**
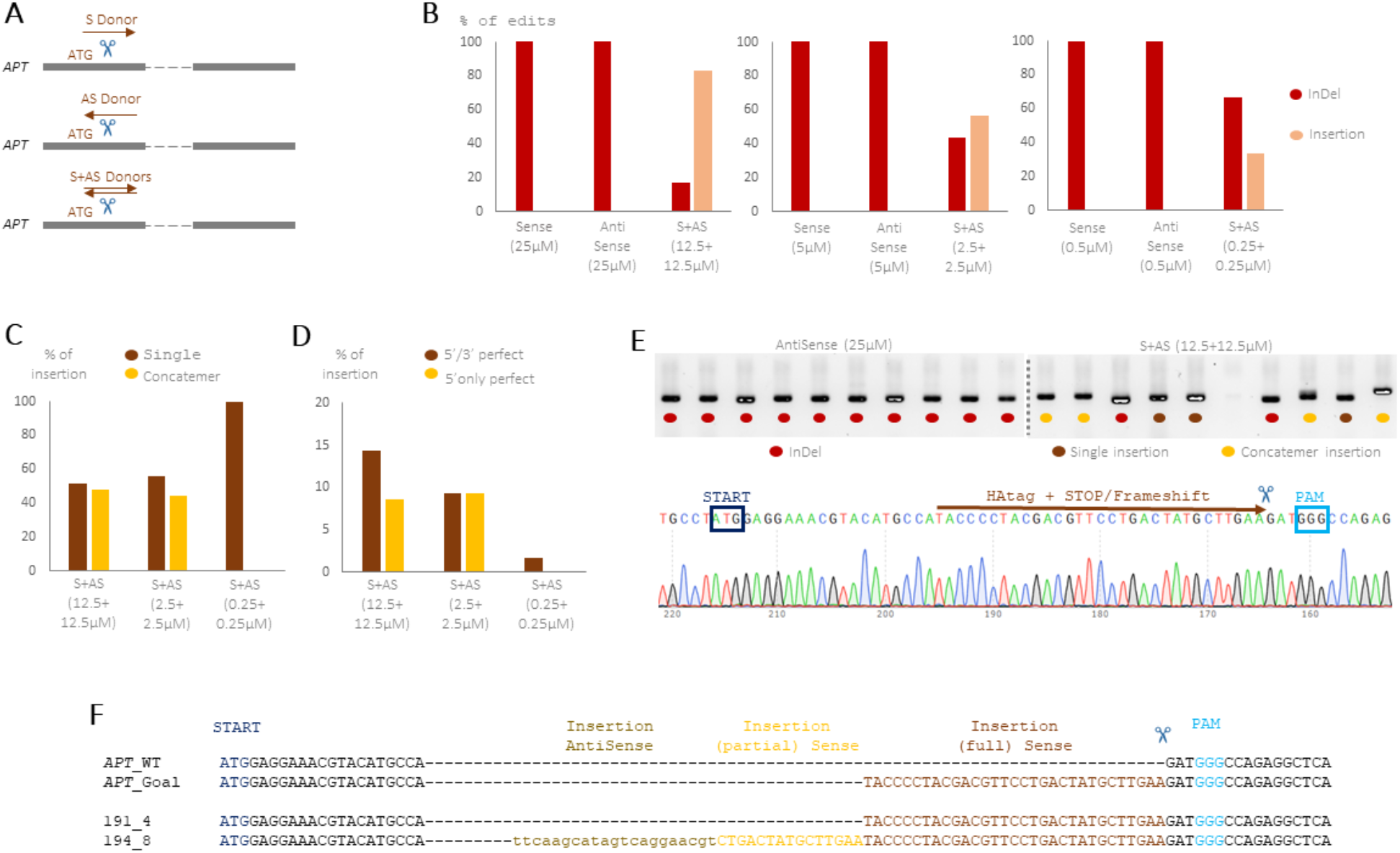
Establishing targeted knock-in at the *APT* locus. (A) Schematic of the KI strategy. The *APT* gene was targeted with Cas9 downstream of the START codon. Donors encode an HAtag followed by a STOP codon and an additional nucleotide to induce a frameshift, and were provided as sense (S) or antisense (AS) ssODNs, or as a mixture of both (S+AS). (B) Distribution of *APT* editing outcomes between InDels and donor insertions at different donor concentrations in the Cas9 transformation mixture (n = 17–58 edited alleles per condition). (C) Distribution of full and partial *APT* donor insertions between single-copy and concatemeric integration events. (D) Percentage of full-length *APT* insertions with perfect 5′ and 3′ donor–genome junctions or a perfect 5′ junction only. (E) Agarose gel of *APT* PCR genotyping, showing insertion-associated band shifts when S and AS donors were provided together. An example of Sanger sequencing confirming a precise donor insertion is shown. (F) Example of an *APT* allele with a perfect insertion and one with concatemer insertions.

Surprisingly, neither ssODN polarity alone produced detectable insertions, whereas combining sense and antisense ssODNs enabled efficient donor integration across a range of donor concentrations, with insertion frequencies generally increasing at higher donor concentrations (**Figure 1B**; **Table S1**). These results suggest that complementary ssODNs may enhance donor stability or availability, or promote their engagement with the DNA repair machinery. Under optimal conditions, up to 83.5% of edited *APT* alleles contained donor-derived insertions, indicating highly efficient capture of exogenous DNA at Cas9-induced DSBs. Increasing donor concentration may increase the local availability of donor molecules at the break; however, insertion efficiency did not scale linearly with concentration. Increasing the donor concentration from 0.25 to 12.5 µM, for example, resulted in only an approximately two-fold increase in the proportion of *APT* alleles carrying an insertion (**Figure 1B**).

Consistent with efficient donor capture, many integration events contained donor concatemers, whose frequency increased with donor concentration (**Figure 1C-F**). No concatemers were detected at the lowest donor concentration, suggesting that donor availability was sufficient for capture at Cas9-induced DSBs but too low to favour the incorporation of multiple donor molecules into a single repair event. We also observed numerous partial donor insertions, suggesting that donor molecules undergo processing or degradation in *Ectocarpus* gametes.

Importantly, many integration events preserved the reading frame at both the 5′ and 3′ donor–genome junctions (**Figure 1D,E**), suggesting that this strategy could be adapted for endogenous protein tagging near the START codon. Other insertions preserved the reading frame only at the 5′ junction. Such events could nevertheless be compatible with C-terminal tagging when targeted immediately upstream of the STOP codon, where the downstream junction lies outside the protein-coding sequence. Together, these results establish ssODN capture at Cas9-induced DSBs as a simple and efficient strategy for endogenous protein tagging in *Ectocarpus*.

### Endogenous protein tagging across multiple loci

We next asked whether the sense + antisense (S+AS) ssODN strategy could be used to tag endogenous genes beyond the *APT* reporter locus. We targeted four genes: *SEC61* (SEC61 translocon subunit), *RAB11* (RAB11 family small GTPase), *MHC* (myosin heavy chain), and *G3BP* (G3BP stress granule assembly factor) (**Figure 2A**). To test the versatility of the approach, we used several epitope tags, including MYC, OLLAS, FLAG and ALFA (**Figure 2B–D**). Editing of each gene of interest (GoI) was performed together with *APT* editing as a co-transformation marker. To favor editing at the GoI, Cas9 RNPs were supplied at a 3:1 ratio of GoI to *APT* guide RNAs, following the principle of co-CRISPR enrichment strategies used in other systems (Akella et al., 2021; Paix et al., 2015).

**Figure 2.**
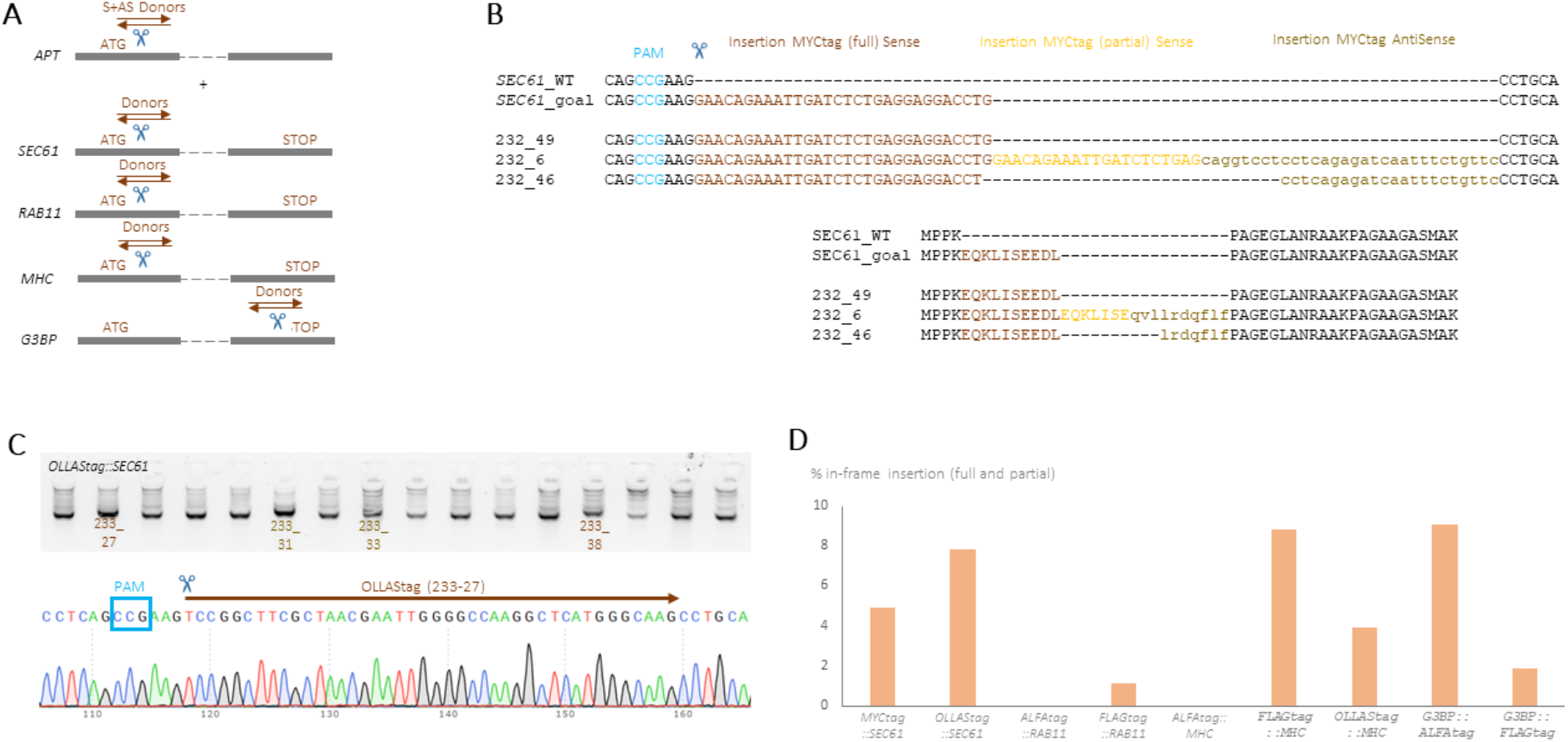
Endogenous protein tagging across multiple loci. (A) Schematic of the endogenous tagging strategy. Genes of interest (GoIs) were targeted either downstream of the START codon (ATG) or upstream of the STOP codon, with simultaneous targeting of *APT* as a co-CRISPR marker. (B) Examples of MYC-tag donor insertions at the *SEC61* locus, illustrating concatemeric, partial and inverted integration events and their predicted translation products. (C) Agarose gel showing PCR genotyping of OLLAS-tag insertions at the *SEC61* locus, including two full-length in-frame insertions (233_27 and 233_38) and two incorrect insertion events (233_31 and 233_33). The Sanger sequencing chromatogram of one full-length in-frame insertion is shown. (D) Percentage of in-frame KI events obtained across several GoIs and epitope tags. Both full-length and partial KIs retaining functional tag sequences are included (n = 0– 4 positive KI lines per experiment).

Using this approach, we obtained targeted insertions at all four loci, with KI frequencies ranging from 1% to 9% among screened transformants (**Figure 2D; Figure S2A; Table S2**). Not all integration events contained the complete donor sequence: some carried partial tags lacking a few codons, whereas others contained fewer tag copies than intended when multimeric donors were used, such as 3×FLAG. For example, one OLLAS-tagged line encoded an 11-amino-acid version of the tag, lacking one N-terminal and two C-terminal residues. In two of nine experiments, the desired tag insertion was not recovered, potentially reflecting the limited number of transformants screened. Because the optimized protocol requires only small numbers of gametes, however, multiple transformations can readily be performed in parallel when a specific KI allele is required. Overall, at least one targeted tag insertion was recovered for each gene tested.

We also found that editing outcomes differed markedly among target loci (**Figure S2B**). When *SEC61* and *RAB11* were targeted close to the START codon, most screened individuals retained the wild-type sequence, and very few, if any, InDels were recovered. This could reflect either relatively inefficient guide RNAs or selection against disruptive mutations in these genes. Strikingly, however, all recovered donor insertions at these loci maintained the coding frame (**Figure S3A**). Moreover, the only *RAB11* InDel recovered removed the original START codon but was accompanied by an upstream SNP that generated a new in-frame ATG, resulting in a predicted RAB11 protein lacking only six amino acids and therefore likely to remain functional (**Figure S3B**). Together, these observations are consistent with *SEC61* and *RAB11* being essential, such that only edits preserving gene function are recovered during gamete and partheno-sporophyte development.

These results also suggest a useful feature of performing KI experiments without prior knowledge of gene essentiality. Because Cas9-induced InDels are generated alongside donor insertions, the spectrum of recovered alleles can provide information about the tolerance of a locus to disruption. A strong enrichment for in-frame insertions, together with the absence of disruptive InDels, may indicate selection against loss-of-function alleles and thus provide evidence for gene essentiality.

This pattern was not observed for *MHC*, which was also targeted downstream of the START codon (**Figure S2B; Figure S3C**). Most sequenced 2-FA-resistant individuals carried edits at the target locus, including numerous insertions and InDels predicted to disrupt the reading frame. This indicates efficient cleavage by the *MHC* guide RNA and suggests that loss of *MHC* function is tolerated under these conditions. In contrast, targeting *G3BP* immediately upstream of the STOP codon yielded a high proportion of wild-type alleles, consistent with relatively low guide-RNA efficiency (**Figure S2B; Figure S3D**). Importantly, the recovered *G3BP* InDels and insertions frequently altered the reading frame, as expected for edits at the C terminus where disruption downstream of the coding sequence is less likely to compromise gene function.

## DISCUSSION

*Ectocarpus* has emerged as a powerful model for investigating the evolution of multicellularity, life-cycle regulation, sex determination and developmental processes in a eukaryotic lineage that evolved complex multicellularity independently of animals and plants (Bringloe et al., 2020; Denoeud et al., 2024). Genomic and transcriptomic resources have enabled increasingly detailed analyses of gene expression and genome evolution, while the recent development of CRISPR-Cas KO approaches has provided the means to directly test gene function (Badis et al., 2021; Martinho et al., 2025). However, the ability to precisely modify endogenous loci has remained a major limitation. Although sophisticated genome engineering and transgenesis approaches are now available in several green algal models (Blomme et al., 2021), these organisms belong to an evolutionarily distant lineage, and comparable tools have remained scarce in brown algae.

Here, we establish targeted genomic insertion in *Ectocarpus* and demonstrate its application for endogenous protein tagging at multiple loci. This represents an important extension of the brown algal genetic toolkit beyond gene disruption, enabling proteins to be studied under their native genomic and regulatory context. We obtained insertions using several peptide tags and at multiple endogenous loci, providing a basis for protein localization, expression and interaction studies without relying on transgene overexpression. In parallel, optimization of PEG-mediated RNP delivery substantially simplified the transformation procedure and improved editing efficiency. Together, these advances establish a practical workflow combining efficient KO and KI within the same experimental framework.

A striking feature of the KI strategy was the strong dependence of donor integration on ssODN configuration. Individual sense or antisense ssODNs produced no detectable insertions, whereas combining complementary ssODNs resulted in efficient donor capture, accounting for up to 83.5% of edited alleles under some conditions. The mechanistic basis of this synergy remains unclear. Complementary ssODNs may anneal before or after cellular entry, generating partially or fully double-stranded substrates that are more stable or more efficiently recognized by DNA repair machinery. Alternatively, the presence of complementary strands could alter donor uptake or processing within the cell. Linear and partially annealed dsDNA molecules can serve as efficient repair substrates in other systems, including *C. elegans* (Dokshin et al., 2018; Ghanta and Mello, 2020; Paix et al., 2015), suggesting that donor structure can strongly influence repair outcomes. Determining the molecular form of the donor that is ultimately incorporated will be important for understanding and further optimizing KI in *Ectocarpus*.

The frequent recovery of concatemeric and partial insertions provides additional insight into donor processing during repair (Smirnov and Battulin, 2021). Increasing donor concentration promoted the incorporation of multiple donor molecules, with up to eight fragments inserted, whereas concatemers were not detected at the lowest concentrations tested. Together with the occurrence of truncated donor insertions, these observations suggest that exogenous DNA undergoes extensive joining and processing in *Ectocarpus* gametes. Importantly, these events occurred in the absence of designed homology either between donor molecules or between the donor and the genomic target, consistent with efficient capture of exogenous DNA through end-joining pathways (Haber, 2000; Mabuchi et al., 2023; Suzuki et al., 2016). Similar chromosomal concatemerization has been reported in zebrafish, although to a lesser extent (Boel et al., 2018). Identifying the repair factors responsible for these outcomes will be important both for understanding DSB repair in brown algae and for improving the precision of genome engineering in this lineage.

Our experiments also illustrate an advantage of combining donor-mediated KI with conventional Cas9 editing. Because InDels are generated alongside donor integrations, a single experiment can simultaneously produce tagged and disruptive alleles. The distribution of editing outcomes may additionally provide information about functional constraint at the targeted locus. For example, the strong enrichment for frame-preserving modifications and absence of disruptive alleles observed at some loci is consistent with selection against loss-of-function mutations. This feature could be particularly valuable in *Ectocarpus*, where the essentiality of most genes is unknown, allowing KI experiments to generate functional alleles while simultaneously revealing whether disruptive mutations are tolerated.

An important advantage of the present strategy is its simplicity. The homology-independent donors used here require minimal locus-specific design and can therefore provide versatile substrates for introducing the same epitope tag at multiple genomic targets. Combined with co-editing of the selectable *APT* locus, this makes endogenous tagging accessible without requiring locus-specific selectable markers or complex donor constructs. The approach should be readily adaptable to additional peptide tags and potentially to other small functional sequences, providing a flexible framework for investigating protein localization and function in *Ectocarpus*.

The current approach nevertheless has limitations. Donor concatemerization and truncation reduce the predictability of integration outcomes and require individual KI lines to be validated by sequencing. Future developments should therefore focus on controlling donor processing and increasing the proportion of precise, single-copy integrations. Donor stabilization or modification may reduce degradation and concatemer formation (Ghanta and Mello, 2020; Yu et al., 2020), while the introduction of homology arms could enable homology-directed repair and more precise sequence replacement (DeWitt et al., 2017; Liao et al., 2024; Mehta and Haber, 2014). At the same time, the simplicity of the current homology-independent strategy remains an important advantage, particularly for routine tagging across multiple loci. The high efficiency of donor capture demonstrated here therefore provides a strong foundation both for immediate applications and for the development of increasingly precise genome engineering approaches in *Ectocarpus*.

Together, our results extend genome engineering in *Ectocarpus* from gene disruption to targeted modification of endogenous loci. The ability to combine KO and KI with endogenous protein tagging creates new opportunities to connect gene function with protein localization and dynamics in a lineage that occupies a key position for comparative studies of eukaryotic development and multicellularity. More broadly, the principles established here provide a foundation for increasingly sophisticated genome engineering across brown algae and potentially other stramenopiles, opening previously inaccessible questions to mechanistic genetic and cell biological analysis.

## ACKNOWLEDGEMENTS

This research was funded by the Max Planck Society and the Gordon and Betty Moore Foundation (SMC). We thank the Culture Facility for assistance with algal cultures and members of the Algal Development and Evolution department for constructive discussions. We also thank the EMBL-Heidelberg PEPcore facility for Cas9 purification.

## MATERIAL AND METHODS

### Algae culture and transformation

*Ectocarpus sp7* were cultured as described before (Coelho et al., 2012). The day of transformation, mature male gametophytes (Ec32) were submitted to artificial low tide in the dark for 3 h. Gamete release was trigger by adding a small volume of ice-cold SW and submitted to strong light. After 20-30 min, gametes were counted using a hemocytometer. Gamete concentration used for transformation ranged from 400 to 8200 gametes/µL, with a range from 5000-7500 gametes/µL for data presented in **Figures 1 and 2**, and no gametes centrifugation was performed to reduce the time of transformation.

Transformation was performed by putting 20 µL of Cas9/Cas12a mix in the center of a 100mm dish, adding 20 (optimized protocol) or 100 µL of gametes (Martinho et al., 2025), and 120 µL of PEG solution. The transformed gametes were kept at 22.5 °C in the dark for 30 min. Next 20 mL of SW+PE (Provasoli’s Enriched, half-strength) was added, and the dishes were kept 14-18 h in the dark at 22.5 °C. The day after, dishes were put in normal culture conditions (14 °C, normal light, normal day) for one day.

The next day, 2-Fluoradenin (2-FA, Sigma-Aldrich, #535087) was added from a 10000X stock (150 mM in DMSO) and cultivated in normal conditions for approximately 2 weeks. Next, SW+PE with 2-FA was changed and cultivated again for approximately 3 weeks. Finally, SW+PE with 2-FA was replaced by SW+PE only and cultivated until 2FA resistant (*APT* mutants) can be recovered and counted. During each media change, the supernatants were kept to isolate *APT* edited partheno-sporophytes which did not attach well to the dishes.

2-FA was used to select for *APT* KO, and in all reported experiments, we did not get false positive with WT *APT* sequence. The large majority of edits resulted in a frame-shift. However, in an instance, we found of an in-frame large deletion (24 bp just downstream the START codon) indicating that the Nt of APT protein is important for its function.

PEG solution was prepared by weighting 6 g of PEG8000 (Fisher BioReagents, #BP233-1) in a 50 mL Falcon tube, and adding 10 mL of SW for a final volume of 15 mL (40% solution), mixed and filtered (0.4 µm).

### Cas9/Cas12a mixes

spyCas9 protein was produced by EMBL Heidelberg PepCore facility as described before (Garriga-Canut et al., 2026; Paix et al., 2019). lbaCas12a was purchased to IDT (Alt-R L.b. Cas12a -Cpf1-Ultra, #10007923). Cas9/Cas12a mixes were prepared the day of transformation, or the day before, by assembling reagents at room temperature, kept on the bench for 15 min, and kept at 4 °C until transformation.

Cas9 mix was assembled by adding the following reagent in this order: 2.5 µL of Cas9 (10-11 µg/µL, in 20 mM HEPES/NaOH pH 7.5, 500mM KCl, 20% glycerol), 2.75 µL of KCl 1M, 0.7 µL of HEPES 0.5M pH7.5, 4 µL total of guide-RNA (100µM), 5.05 µL H2O, 5 µL of total donor or 5 µL of H2O if no donors were used.

Cas12a mix was assembled by adding the following reagent in this order: 2 µL of Cas12a (10 µg/µL, in IDT buffer), 4 µL of total guide-RNA (100 µM), 2 µL of 3.1r 10X buffer (NEB, #B7203S), 6 µL of IDTE buffer pH7.5 (IDT, #11-05-01-15), 1 µL H2O, 5 µL of total donor or 5 µL of H2O if no donors were used.

If only *APT* was targeted, 4 µL of corresponding guide-RNA was used. If a GoI was also targeted, *APT* was used as a selection marker, and 1 µL of *APT* guide-RNA was used together with 3 µL of guide-RNA targeting the GoI.

Cas9 guide-RNAs were purchased custom made with Sigma-Aldrich (sgRNA, standard purification, no modification) and reconstituted in H2O, stored at -80 °C. *APT* Cas12a guide-RNA was purchased custom made with IDT (standard desalting) and reconstituted in IDTE buffer pH7.5, stored at -80 °C. Guide-RNA were manually selected to be as close possible to the START and STOP codon, with few potential off-targets (determined using CRISPOR) (Concordet and Haeussler, 2018), and having a GC content between 30% and 80% when possible (Doench et al., 2014; Gagnon et al., 2014). Available gene model and genomic sequence resources were used for design (Barrera-Redondo et al., 2025; Denoeud et al., 2024).

ssODN donors and primers were purchased custom made with Sigma-Aldrich (standard desalting) at 100 µM in H2O and stored at -20 °C. APm233/APm234 were purchased with 5’biotin. When ssODN were used at high concentration as for targeting the GoI, 2.5 µL of sense donor (100 µM) and 2.5 µL of antisense donor (100 µM) were used in the 20 µL Cas9 mix.

### Genotyping

Individual 2-FA resistant were transferred in 24-wells plates containing SW+PE. Small pieces of each individual were cut, lysed, and vortexed in 96-wells plates. PCR genotyped was performed using Phire Plant Direct PCR Master Mix (Thermo Scientific, # F160L). Lysing buffer was 40 µL of H2O and 40 µL Phire lysis buffer, with stainless steel beads (Carl Roth, #P329.1). PCR mixes were: 10 µL of 2X Phire Plant Direct PCR Master Mix, 0.1 µL Forw-primer (100 µM), 0.1 µL Rev-primer (100 µM), 9.3 µL H2O, 0.5 µL DNA lysis. PCR conditions were: 98 °C for 5 min; 35x of 98 °C for 5 sec / annealing (60 to 72 °C) for 5 sec / 72 °C for 40 sec; 72 °C for 1 min. PCR products were analyzed in mini-agarose gels. Sanger sequencing were performed by Azenta (Leipzig, Germany) using PCR product purified with ExoSAP-IT PCR Product Cleanup Reagent (Applied Biosystems, #78201.1.ML).

Some insertions were difficult to Sanger sequence due to the presence of concatemer insertions, and the genotyping PCR were purified using Minelute PCR cleanup (Qiagen, #28004) and Nanopore sequenced using Azenta PCR-EZ service

Positive KI individuals were propagated by excising a small fragment of tissue and allowing it to grow in fresh wells or culture dishes. The resulting lines were subsequently re-genotyped at both the gene of interest (GoI) and the *APT* locus to confirm the edits. Validated lines will be made available upon request. Guide RNA and ssODN sequences, genotyping primers, and the corresponding available lines are listed in **Tables S3 and S4**.

## SUPPLEMENTAL FIGURES

**Figure S1.**
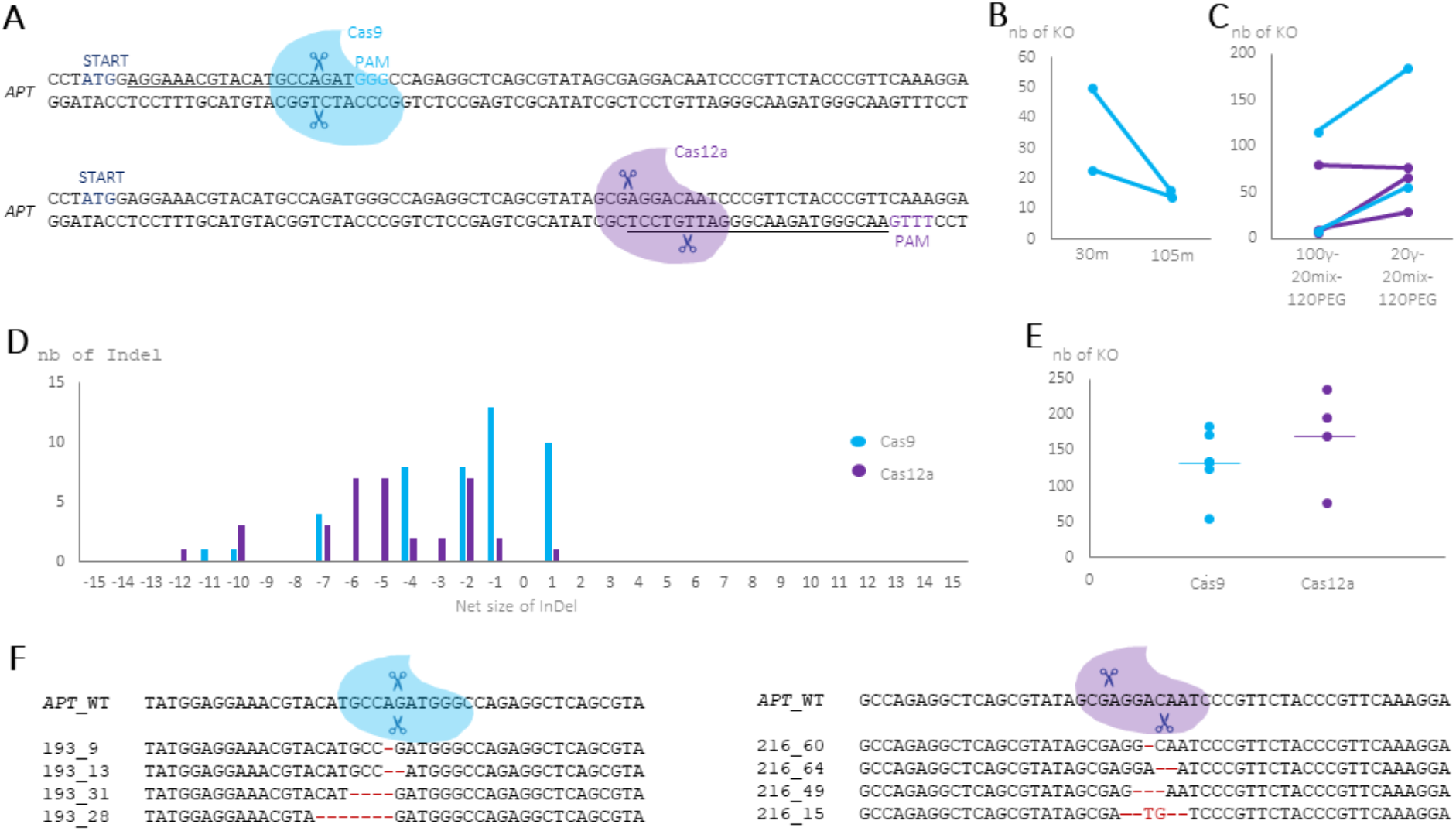
Optimization of transformation and editing conditions. (A) *APT* sequences targeted by Cas9 (blue) and Cas12a (purple), with the guide-RNA spacer sequences underlined. Cas9 generates predominantly blunt-ended double-strand breaks, whereas Cas12a generates staggered cuts with 5′ overhangs. (B) Number of *APT* edits recovered when the same Cas9 RNP mixture was used to transform aliquots from the same batch of gametes at 30 or 105 min after release. (C) Number of *APT* edits recovered when the same Cas9 or Cas12a RNP mixture was used to transform aliquots from the same batch containing either 20 µL or 100 µL of gametes (γ), together with 20 µL of RNP mixture and 120 µL of PEG solution. (D) Distribution of *APT* InDel sizes generated by Cas9 and Cas12a. Only InDels within a 30-bp window are shown. Among 49 Cas9-edited alleles, the largest deletion was 46 bp and the mean net InDel size was −3 bp. Among 41 Cas12a-edited alleles, the largest deletion was 61 bp and the mean net InDel size was −9 bp. (E) Number of *APT* edits recovered with Cas9 and Cas12a under optimized conditions. Because Cas9 and Cas12a target different sequences, their efficiencies cannot be directly compared; however, both nucleases generated large numbers of edited individuals when used at high RNP concentrations. Horizontal bars indicate mean values (135 for Cas9 and 170 for Cas12a). (F) Representative *APT* InDels generated by Cas9 and Cas12a.

**Figure S2.**
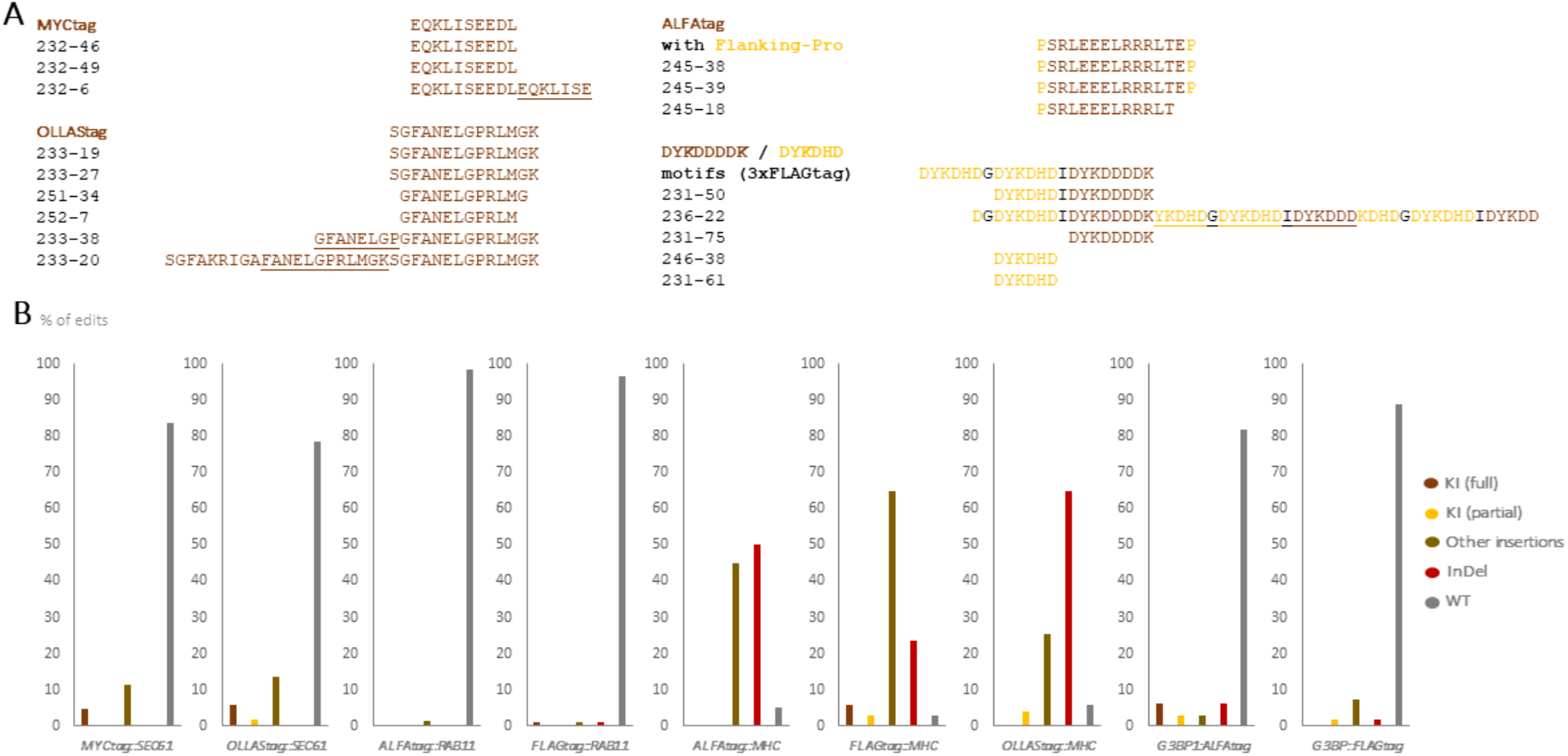
Tag insertions and editing outcomes across endogenous target genes. (A) In-frame insertions obtained with different epitope tags. For concatemeric insertions, individual tag repeats are underlined. (B) Distribution of editing outcomes across different genes and tag donors. Partial KI denotes donor insertions in which the tag sequence lacks one or more codons (n = 32–90 genotyped individuals per experiment).

**Figure S3.**
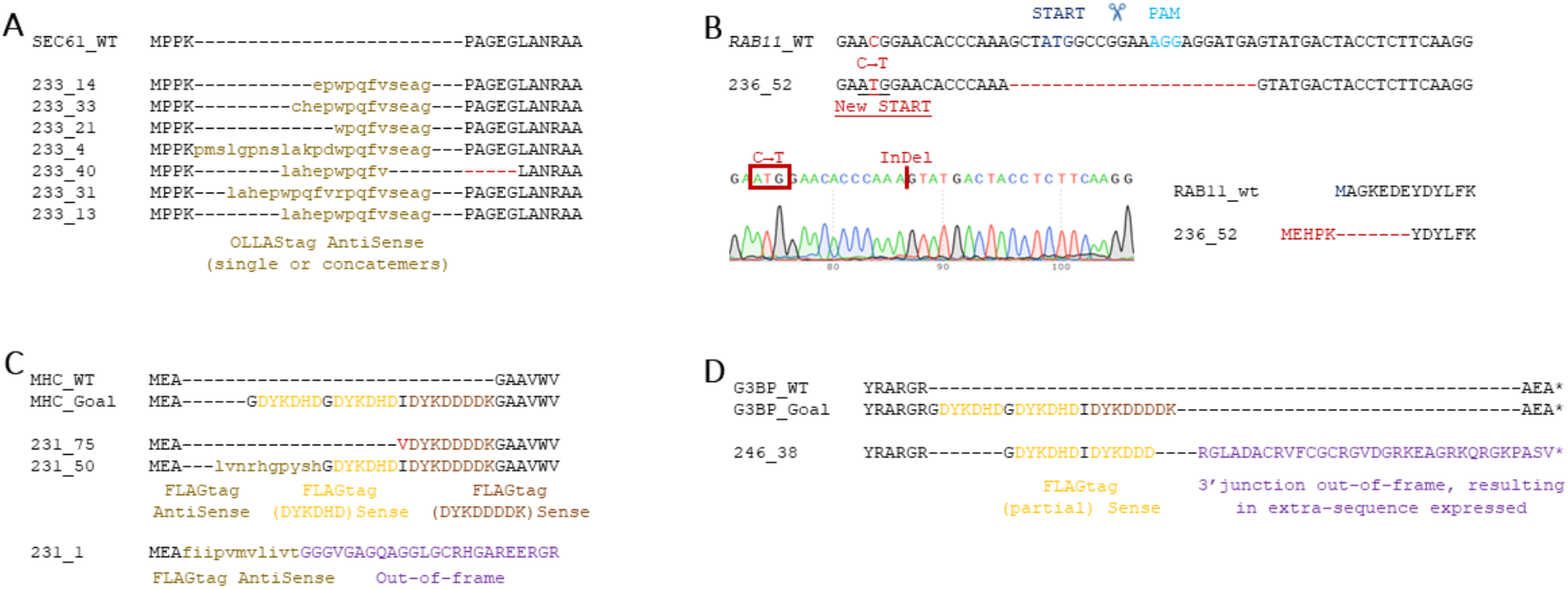
Representative editing outcomes at endogenous loci. (A) Representative non-desired insertions at the N-terminus of SEC61, all of which preserve the reading frame. (B) Sequence of the only InDel recovered at the *RAB11* locus, showing a deletion encompassing the original START codon and an upstream SNP generating a new ATG in frame with the *RAB11* coding sequence. (C) Representative insertions at the N-terminus of MHC. (D) Example of a *G3BP* insertion near the STOP codon in which the 5′ junction is in-frame but the 3′ junction is not, resulting in the generation of a new downstream STOP codon.

## SUPPLEMENTARY TABLES

**Table S1**. Detailed *APT* editing outcomes using ssODN donors.

**Table S2**. Detailed editing outcomes at genes of interest (GoIs) using ssODN donors.

**Table S3**. Validated KI lines generated in this study.

**Table S4**. ssODN donor, guide RNA and PCR/genotyping primer sequences.

